# Biochemical and structural studies of target lectin SapL1 from the emerging opportunistic microfungus *Scedosporium apiospermum*

**DOI:** 10.1101/2020.11.10.376251

**Authors:** Dania Martínez-Alarcón, Viviane Balloy, Jean-Philippe Bouchara, Roland J. Pieters, Annabelle Varrot

## Abstract

*Scedosporium apiospermum* is an emerging opportunistic fungal pathogen responsible for life-threatening infections in humans. Host-pathogen interactions often implicate lectins that have become therapeutic targets for the development of carbohydrate mimics for antiadhesive therapy. Here, we present the first report on the identification and characterization of a lectin from *S. apiospermum* named SapL1. SapL1 was found using bioinformatics as a homolog to the conidial surface lectin FleA from *Aspergillus fumigatus* known to play a role in the adhesion to host glycoconjugates present in human lung epithelium. In our strategy to obtain recombinant SapL1, we discovered the importance of osmolytes to achieve its expression in soluble form in bacteria. Analysis of glycan arrays indicates specificity for fucosylated oligosaccharides as expected. Submicromolar affinity was measured for fucose using isothermal titration calorimetry. We solved SapL1 crystal structure in complex with α-methyl-L-fucoside and analyzed its structural basis for fucose binding. We finally demonstrated that SapL1 binds to bronchial epithelial cells in a fucose-dependent manner. The information gathered here will contribute to the design and development of glycodrugs targeting SapL1.

## Introduction

During the last decades, an increased incidence of invasive infections, especially in immunosuppressed patients, has been caused by previously rare fungal pathogens such species from the *Scedosporium* genus^1,2^. This genus comprises more than ten worldwide distributed soil saprophyte species whose taxonomy has been previously described based on molecular phylogenetic^3,4^. *Scedosporium* species may lead to bronchitis and allergic bronchopulmonary mycoses as well as severe disseminated infections through inhalation of conidia. They rank second among the filamentous fungi colonizing the respiratory tract of cystic fibrosis (CF) patients, after *Aspergillus fumigatus*^4^. Together with *S. boydii* and *S. aurantiacum, S. apiospermum* is the most common *Scedosporium* species able to chronically colonize the lungs of CF patients^4,5^. Because of their propensity to disseminate in case of immune deficiency, this fungal colonization of the airways is considered in some centers as a contraindication to lung transplantation that is the ultimate treatment in CF^6^. Besides, infections by *Scedosporium* species have also been reported in immunocompetent patients^7^.

The treatment of *Scedosporium* infections is challenging because its efficacy depends on the biological diagnosis which is a time-consuming process. The common co-colonization by other fungi or bacteria may lead to false negative results, especially from respiratory secretions of patients with CF^4^. Furthermore, *Scedosporium* species display a primary resistance to most current antifungals such as amphotericin B or first-generation triazole drugs as fluconazole or itraconazole and exhibit a limited susceptibility to the newest generations of antifungal drugs, *i.e.* echinocandins and voriconazole^4,8^. Nowadays, the first-line treatment involves combination therapy but following common recurrences and even without interruption of treatment, the recovery rates are poor and mortality remains over 65% while it is almost 100% when dissemination occurs, reason why it has aroused special interest^8–10^.

*Scedosporium* infections begin with conidial adherence to tissues, followed by germination and hyphal elongation^11^. The adherence allows it to avoid cleansing mechanisms aimed to eradicate the invading pathogens and represents the initial step towards infection^11–14^. Conidial adhesion is mediated by cell surface molecules (CSM), including carbohydrates where some of the most important described to date for *Scedosporium* species include peptidorhamnomannans (PRMs)^15–17^, α-glucan^18^, melanin^19^, ceramide monohexosides^17,20^, N-acetyl-D-glucosamine-containing molecules^21^ and mannose/glucose-rich glycoconjugates^17^. Their presence and/or abundance on the cell surface vary according to the stage of development and is of great relevance to understand fungal pathobiology^17^. The carbohydrates binding proteins known as lectins also act as CSMs and were shown to have an essential role during pathogenesis in the host recognition and adhesion process^22^. They became drug targets for the development of carbohydrates related molecules as antiadhesives drugs^12–14,23–26^. The anti-adhesive therapy approach is particularly promising since it does not kill the pathogen nor arrest its cell cycle. Consequently, resistance frequencies are rare^13,23^ and expected side effects in non-target tissues are lower than those caused by conventional antifungal compounds. This may allow its implementation as a prophylactic therapy for immunocompromised patients^14^.

Due to the emerging character of *S. apiospermum*, there is very limited information on its mechanisms of recognition and anchoring to the host. Furthermore, lectins from this microorganism have not been characterized to date, what hinders the development of an anti-adhesive therapy. Conversely, other filamentous fungal pathogens were investigated leading to the identification and characterization of their host binding modes. For example, in *A. fumigatus,* which is a saprophytic mold also responsible for bronchopulmonary infections in receptive hosts, the lectin FleA (or AFL) was identified and revealed a role in host-pathogen interactions^27^. FleA is a six-bladed β-propeller homodimer located on the conidial surface that recognizes human blood group antigens and mediates *A. fumigatus* binding to airway mucins and macrophages glycoproteins in a fucose-dependent manner^28^. In healthy individuals, this anchorage is critical for the mucociliary clearance process and the macrophagy; in fact, it has been described that fleA-deficient (*ΔfleA*) conidia are even more pathogenic than wild type (WT) conidia, both in healthy and chemically immunocompromised mice^28,29^. However, CF patients represent a very particular scenario because the mucus in their lungs is thicker, in relation with mucin overproduction and its high content in calcium ions, which modulates the supramolecular organization of mucin MUC5B by protein cross-linking^29,30^. This contributes to the suboptimal transport properties of mucus and compromises the pathogen clearance mechanisms^29^. Furthermore, the aberrant glycosylation in CF patients causes, among other things, an increase in the abundance of sialyl-Lewis X and Lewis X determinants in lung mucins^31,32^, which is translated as an increase in the fucose content. Therefore, in this context, the FleA (and homologous proteins) anchoring to the mucus layer plays an essential role in the colonization of the CF lungs by *A. fumigatus*.

Here, we have used the recently sequenced genome of *S. apiospermum*^33^ to identify a putative homologue of FleA that we have called SapL1 for *Scedosporium apiospermum* Lectin 1. The present report comprises SapL1 identification, its expression in soluble form in bacteria, an analysis of the fine specificity and affinity of the recombinant protein, as well as its structural characterization by X–ray crystallography and fucose-dependent binding to bronchial epithelial cells by fluorescence microscopy.

## Results

### Production and purification of SapL1

The hypothetical protein XP_016640003.1 (EMBL accession number), encoded by SAPIO_CDS9261 and from now on referred as SapL1, was identified through data mining using the FleA (pdb entry 4D4U^34^) sequence as bait into the genome of the reference strain *S. apiospermum* IHEM 14462^33^. During the sequence analysis, we found that the first 74 amino acids of the putative protein (Uniprot A0A084FYP2) represent a very disordered region that is not present in any other related protein. We suspected that this peptide could be due to an error during the genome annotation. To verify it, we tried to get the gene from total mRNAs extracted from mycelium by 5’ Rapid Amplification of cDNA Ends (5’ RACE) using the following primers GSP1-SapL1 5’-TTAAGCAGGGGGCAGAACAGC-3’ and GSP2-SapL1 5’-CCAACGACCCAGCCAGAGTTCC-3’ but we were unsuccessful. This could be due to the fact that the gene was not expressed in the conditions tested. We did not have access to total mRNAs from other morphological stages of the microfungus. Thus, we decided to focus on the identified carbohydrate recognition domain (CRD) of SapL1 starting from methionine-75 (Fig. S1). The SapL1 coding sequence (75-369) was fused to an N-terminal 6xHis tag cleavable by the Tobacco Etch virus protease (TEV) under regulation of *trc* and *T7* promoters into pProNde and pET-TEV vectors, respectively (Fig. 1A). Expression was performed in *Escherichia coli* and purification was carried out using immobilized metal affinity chromatography (IMAC). Unfortunately, the original expression yield with both vectors (~0.35 mg·L^−1^ of culture) was too low to proceed with characterization studies. Therefore, we explored new alternatives to enhance the expression. First, the thioredoxin protein (Trx) as well as 6-His tag and TEV cleavage site were fused at the N-terminus of SapL1 by subcloning into the pET32-TEV vector (Fig. 1A). This strategy substantially increased the SapL1 production yield but most of the protein remained insoluble as part of inclusion bodies (Fig. S2).

**Figure 1.**
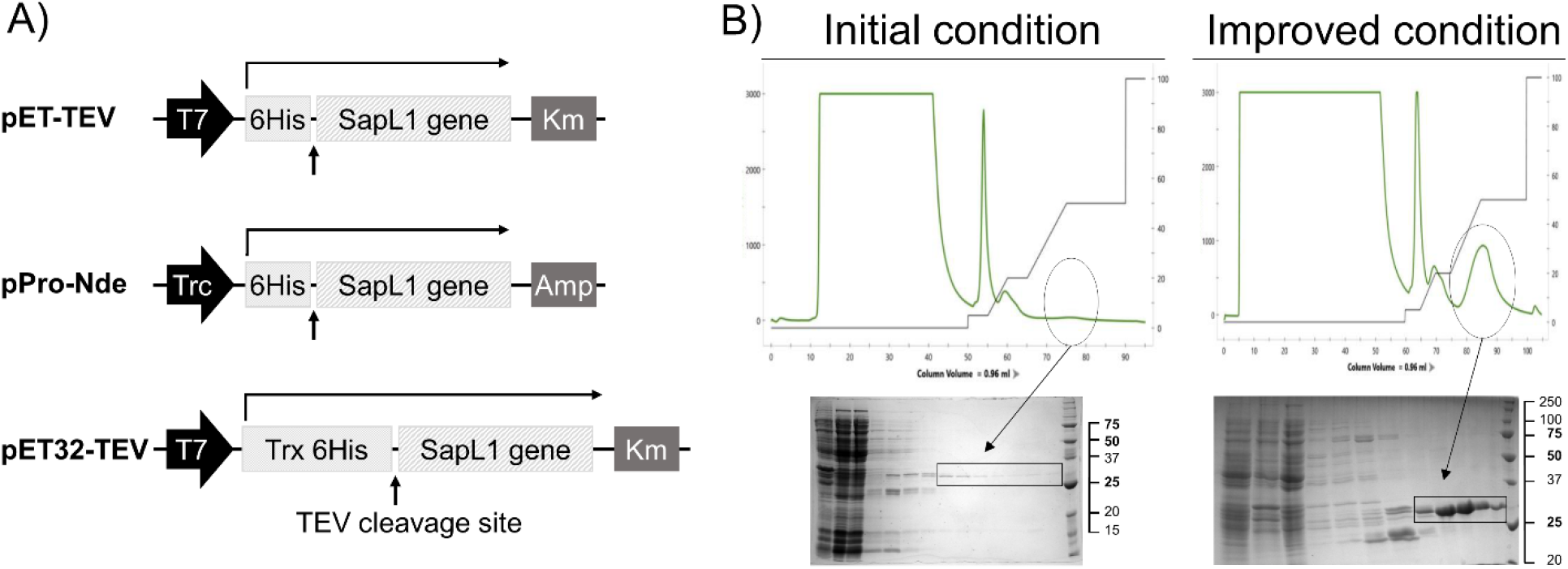
SapL1 production. A) Schematic representation of the genetic constructs used for the expression of SapL1. B) Representative chromatograms of SapL1 purification before and after the process with their respective profile on 15% SDS-polyacrylamide gels (insoluble fraction, soluble fraction, flow through, washing and elution and molecular weight marker, from left to right respectively). The fractions containing SapL1 are delimited.

A wide range of expression conditions were subsequently assayed to achieve sufficient soluble expression by modification of various parameters such as growth temperature, host strain, inducer concentration, optical density of the culture at induction, culture duration, etc. Sixty-nine different sets of parameters were assayed (Table S1) and it was possible to improve the yield up to 4 mg·L^−1^ (Fig. 1B). The best set of conditions for SapL1 expression was using *E. coli* strain TRX, pProNde vector, LB medium, growth at 37°C and 160 rpm until OD_600_ = 0.4, before a switch of the temperature to 16°C and overnight induction at OD 0.8 with 0.05 mM IPTG and 1% L-rhamnose (Rh).

### Rhamnose influence on SapL1 solubility

Interestingly, during the optimization of the expression conditions, we found that rhamnose (Rh) plays an essential role in the solubility and stability of SapL1. Therefore, we performed a new set of experiments to demonstrate the correlation between the amount of protein recovered after purification and the Rh concentration in the media. For this, we investigated SapL1 expression under the previously described conditions modified by supplementation of the culture medium with different concentrations of Rh (0.1%, 0.15%, 0.2%, 0.5% and 1%). Purification parameters were set to obtain high purity product and kept identical for all experiments. Figure 2 shows the chromatograms obtained for representative concentrations accompanied by their respective SDS-PAGE profile. To quantify SapL1 expression in these experiments, we integrated the area under peaks corresponding to the protein (Fig. 2, black bars). Thus, we confirmed that SapL1 recovery is directly proportional to the Rh concentration in the culture medium. Then, the experiment was repeated at three different concentrations (0.1%, 0.15%, 0.2%) using 20°C as the induction temperature, and production behavior was similar to that found at 16°C, but with lower performance.

**Figure 2.**
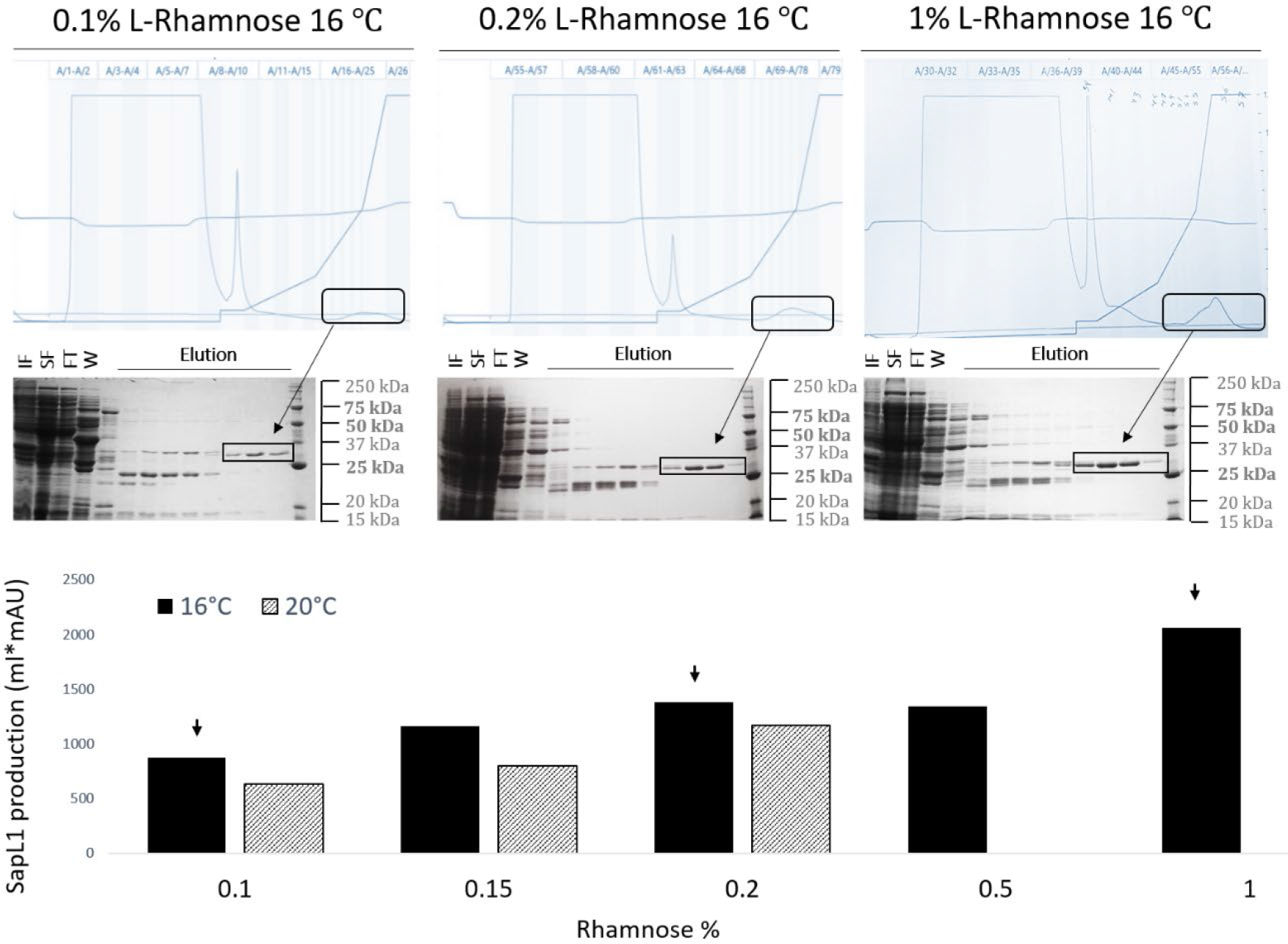
Rhamnose influence on SapL1 solubility. A) SapL1 purification chromatograms at 0.1%, 0.2%, and 1% rhamnose with their corresponding SDS-PAGE profile. IF: Insoluble fraction, SF: Soluble fraction, FT: Flow through, W: wash and elution, from left to right respectively. Black rectangles indicate the elution peaks and elution fractions containing SapL1 on SDS-polyacrylamide gels. B) Numerical values derived from the integration of areas under the peaks corresponding to SapL1 for each experiment. Black and striped bars represent the results for experiments performed at 16°C and 20°C, respectively.

Additional experiments also showed that SapL1 expression can be induced at high concentration of rhamnose in a dose-dependent manner, even without addition of IPTG when the pProNde vector is used (data not shown).

### Biochemical characterization

We assayed the thermal stability of SapL1 in 26 different buffers in a pH range of 5 to 10 through Thermal Shift Assay (TSA). The most suitable condition for this protein was MES buffer 100 mM pH 6.5, where a single denaturing event at T_m_ of 55°C was observed (Fig. S3). Then, to estimate the molecular size and oligomerization state of the native protein, we performed size exclusion chromatography using an ENrich™ SEC 70 column (Bio-Rad) and 20 mM MES, 100 mM NaCl, pH 6.5 as mobile phase. However, the protein displayed strong non-specific interactions with the matrix of the column and SapL1 could not be eluted even using 5 M of NaCl. Interestingly, it could be recovered when the buffer was supplemented with 20 mM α-methyl fucoside or L-rhamnose, evidencing similar effects on SapL1 elution for both sugars. Due to the impossibility to estimate the molecular weight of SapL1 by size exclusion chromatography on this resin, we performed measurements in solution using Dynamic Light Scattering (DLS). We obtained a monodisperse peak corresponding to a protein of 72 ± 29.4 kDa corroborating that SapL1 forms dimers as FleA and other proteins of this family (monomer MW: 40 kDa, data not shown)^27^. The range of the standard deviation also suggested an ellipsoidal shape, which is characteristic for the dimers in this lectin family^27,35^.

### Carbohydrate binding properties

A hemagglutination assay showed that recombinant SapL1 agglutinated rabbit red blood cells at 0.97 μg·mL^−1^ (Fig. 3A). It confirms that the recombinant lectin is active and that its heterologous production in *E. coli* did not alter its hemagglutinating properties.

**Figure 3.**
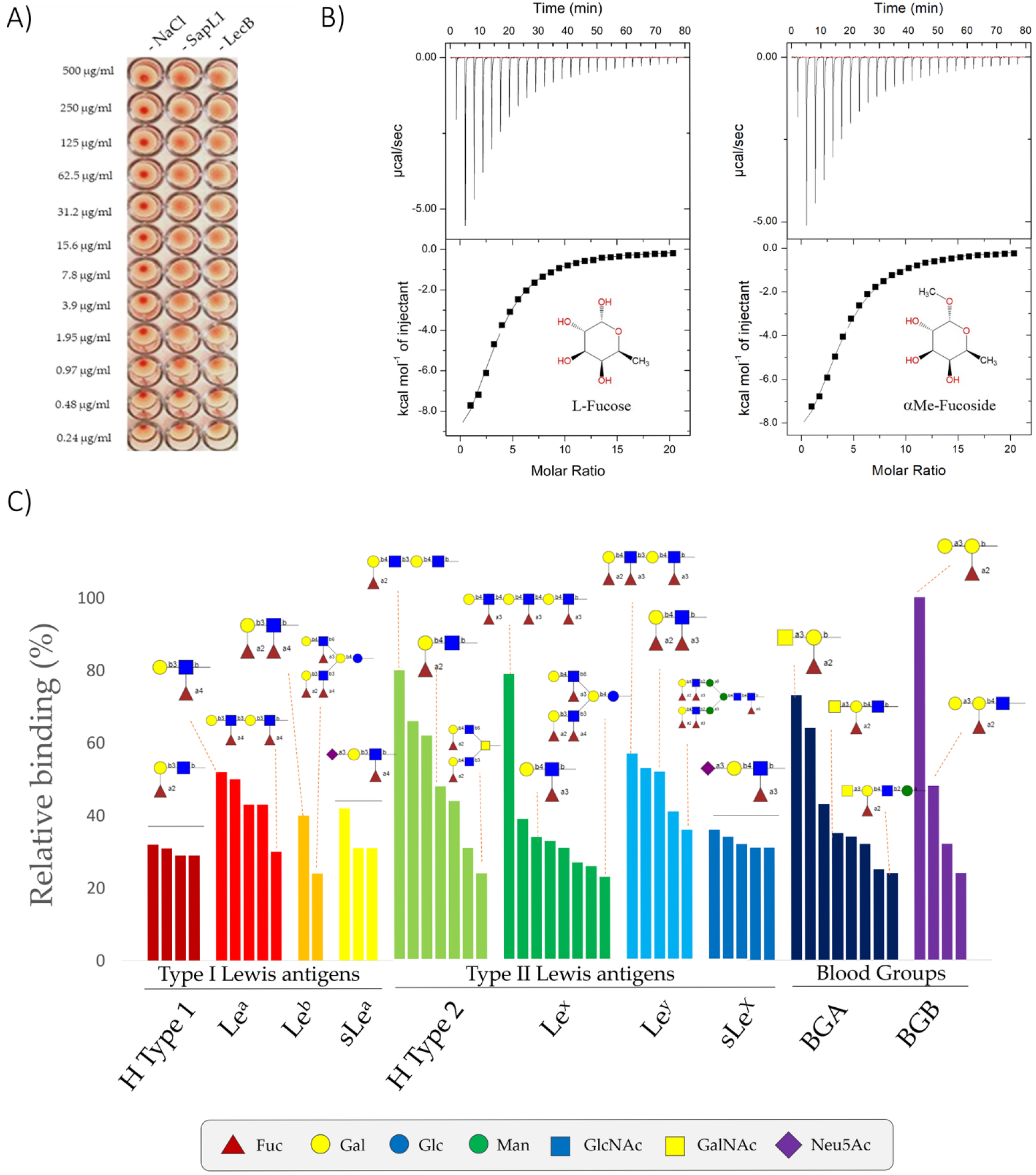
SapL1 carbohydrate binding properties. A) Hemagglutination assay of SapL1 on fresh rabbit erythrocytes. Negative and positive controls consist of 150 mM NaCl and the lectin LecB from *Pseudomonas aeruginosa*, respectively. B) Titration of SapL1 with L-fucose and α-methyl-fucoside with the thermogram and the integration displayed at the top and bottom, respectively. C) Analysis of the interactions of SapL1 with blood group epitopes. The graph shows the relative binding of SapL1 to glycans containing Lewis and ABH blood group antigens from the 90 hits identified as binders.

To identify the potential ligands of SapL1 on epithelial cell surfaces, we submitted SapL1 to the glycan array version 5.4 of the Consortium for Functional Glycomics (USA) consisting of 585 mammalian glycans. It was labelled with Fluorescein Isothiocyanate (FITC) in a molar ratio of 0.426 and its binding properties were analysed at two different concentrations (5 and 50 μg·mL^−1^). As expected from its homology with FleA, SapL1 recognizes fucosylated oligosaccharides independently of the fucose linkage. The α1,2 and α1,3/4 linked fucosides displayed the highest affinity whilst the lowest was seen with the α1,6 linked ones. The weakest interactions with fucosylated compounds were reported for branched oligosaccharides (Figs. 3C and S4).

The affinity of SapL1 for L-fucose and α-methyl-fucoside was determined by Isothermal Titration Calorimetry (ITC) and the K_d_ was found to be 225 ± 1.52 μM and 190 ± 1.44 μM, respectively with stoichiometry fixed to 1 since the measurements were done in the presence of an excess of ligand (Fig. 3B). These values are in agreement with the affinity constant (around 110 μM) reported for FleA for α-methyl-fucoside^36^. No binding interaction was observed for SapL1 with rhamnose by ITC (data not shown).

Analysis of the glycans constituting blood group determinants revealed that SapL1 binds to all epitopes with a preference for H type 2 (Fucα1-2Galβ1-4GlcNAcβ) then Lewis^a^ (Galβ1-3(Fucα1-4)GlcNAcβ) and Lewis^x^ (Galβ1-4(Fucα1-3)GlcNAcβ). However, most of the recognized branched oligosaccharides contained the core fucose Fucα1-6. Epitopes with two fucose units, such as Lewis^b^ and fucosylated polylactosamine, were also well recognized. Addition of a galactose or a GalNAc as in blood group B or A antigens did not impair Fucα1-2 recognition (Fig. 3C). Spacers used to join the carbohydrates to the chip also display a strong influence on binding. It is remarkable that 70% of the 90 positive binders contained either the spacer Sp0 (CH_2_CH_2_NH_2_) or Sp8 (CH_2_CH_2_CH_2_NH_2_). Those spacers also display a strong influence on binding, especially for small glycans such as Galα1-3(Fucα1-2)Galβ1-4(Fucα1-3)GlcNAcβ which was recognized when attached to Sp0 but not to Sp8. This may be due to a steric hindrance caused by the modification of carbohydrate presentation on the surface of the chip.

### Overall structure of SapL1

SapL1 was co-crystallized with α-methyl-fucoside and the structure of the complex was solved by molecular replacement at 2.3 Å resolution in the P2_1_ space group using the coordinates of FleA (PDB code 4D4U^34^) as the search model. The asymmetric unit contained two monomers, assembled as a dimer with all 295 amino acids visible apart of the N-terminal methionine. See data collection and refinement statistics in Table 1. SapL1 folds into the canonical six-bladed β-propeller with six-binding sites at the interface between blades typical for this family of lectin (Fig. 4). A fucose moiety was found in 5 and 4 of the six binding pockets of chains A and B, respectively while glycerol originating from the crystallizing solution was found in the other binding sites.

**Table 1.** Data collection and refinement statistics. *Values in parentheses are for the outer shell

| Data collection |  |  |
| --- | --- | --- |
| Beamline | SOLEIL Proxima 1 |  |
| Wavelength (Å) | 0.97857 |  |
| Space group | P2<sub>1</sub> |  |
| Unit cell dimensions a, b, c (Å), $\alpha$, $\beta$, $\gamma$ (°) | 76.06, 45.66, 83.48, 90, 105.05, 90 | |
| Resolution (Å) | 40.0 – 2.4 (2.46-2.4) |  |
| *R*<sub>merge</sub> | 0.085 (0.428) |  |
| *R*<sub>pim</sub> | 0.07 (0.356) |  |
| Mean *I*/$\sigma$ (*I*) | 5.3 (1.6) | |
| Completeness (%) | 98.2 (98.4) |  |
| Multiplicity | 2.7 (2.7) |  |
| CC<sub>1/2</sub> | 0.991 (0.7) |  |
| No. reflections /No. Unique reflections | 58305/ 21570 |  |
| Refinement |  |  |
| Resolution (Å) | 40.0-2.4 |  |
| No. of reflections in working set / Free set | 21561 / 1089 |  |
| *R* work/ *R* free | 17.5 / 24.0 |  |
| R.m.s Bond lengths (Å) | 0.017 |  |
| Rmsd Bond angles (°) | 1.94 |  |
| Rmsd Chiral (Å<sup>3</sup>) | 0.087 |  |
| No. atoms / Bfac (Å<sup>2</sup>) | Chain A | Chain B |
| Protein | 2222 / 32.6 | 2220 / 40.8 |
| Ligand and heterogen | 89 / 31.6 | 66 / 36.9 |
| Waters | 111 / 31.0 | 69 / 33.2 |
| Ramachandran Allowed / Favored/ Outliers (%) | 96 / 4 / 0 | 95 / 4 / 1 |
| PDB Code | 6TRV |  |

**Figure 4:**
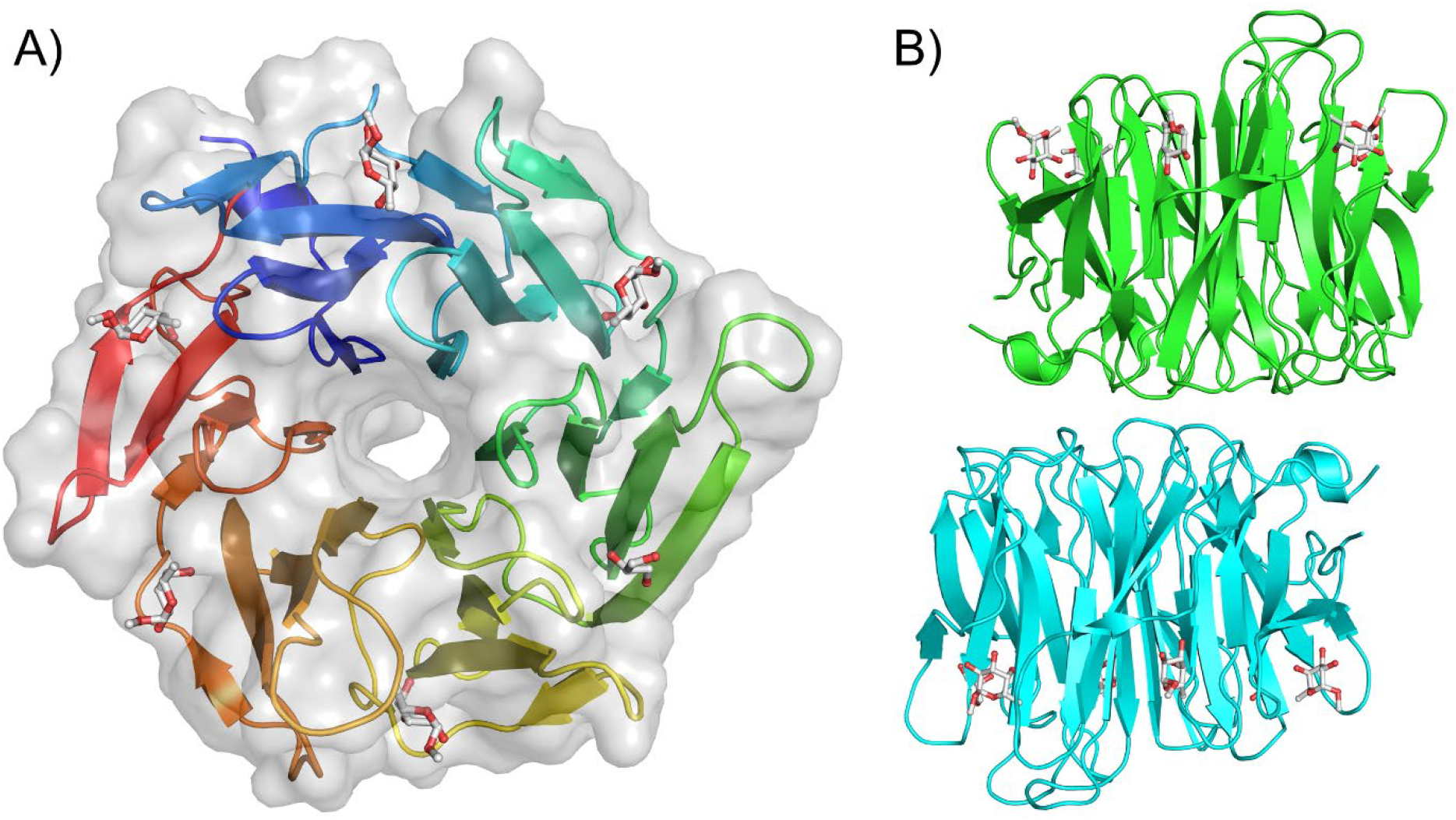
SapL1 overall structure. A) Surface and cartoon representation of SapL1 monomer colored from blue (*N*-terminal end) to red (*C*-terminal end). BP: Binding pockets. B) Representation of SapL1 dimer colored by monomer with ligand depicted in sticks.

The overall fold of SapL1 and FleA as well as the overall dimer are very similar with a rsmd of 1.2 and 1.26 Å, respectively. They share 43% of sequence identity and both proteins present the same distribution of β-strands except for the lack of the last blade of strand 4 in SapL1, and the external face of blade 5, in which FleA displays an elongatedβ-strand (β22) that is split in two in SapL1 (β22/23). It is remarkable that the structure of the first 3 blades is highly conserved in both proteins, while the second half (blades 3-6) displays the largest discrepancies. Furthermore, FleA also presents two additional small α-helices (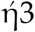 and 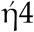), located in the loops between sheets that serve as connectors for blades (2/3 and 5/6, respectively, Fig. 5). Similar conclusions can be drawn when comparing SapL1 to its homologue AOL in *Aspergillus oryzae^37^*.

**Figure 5:**
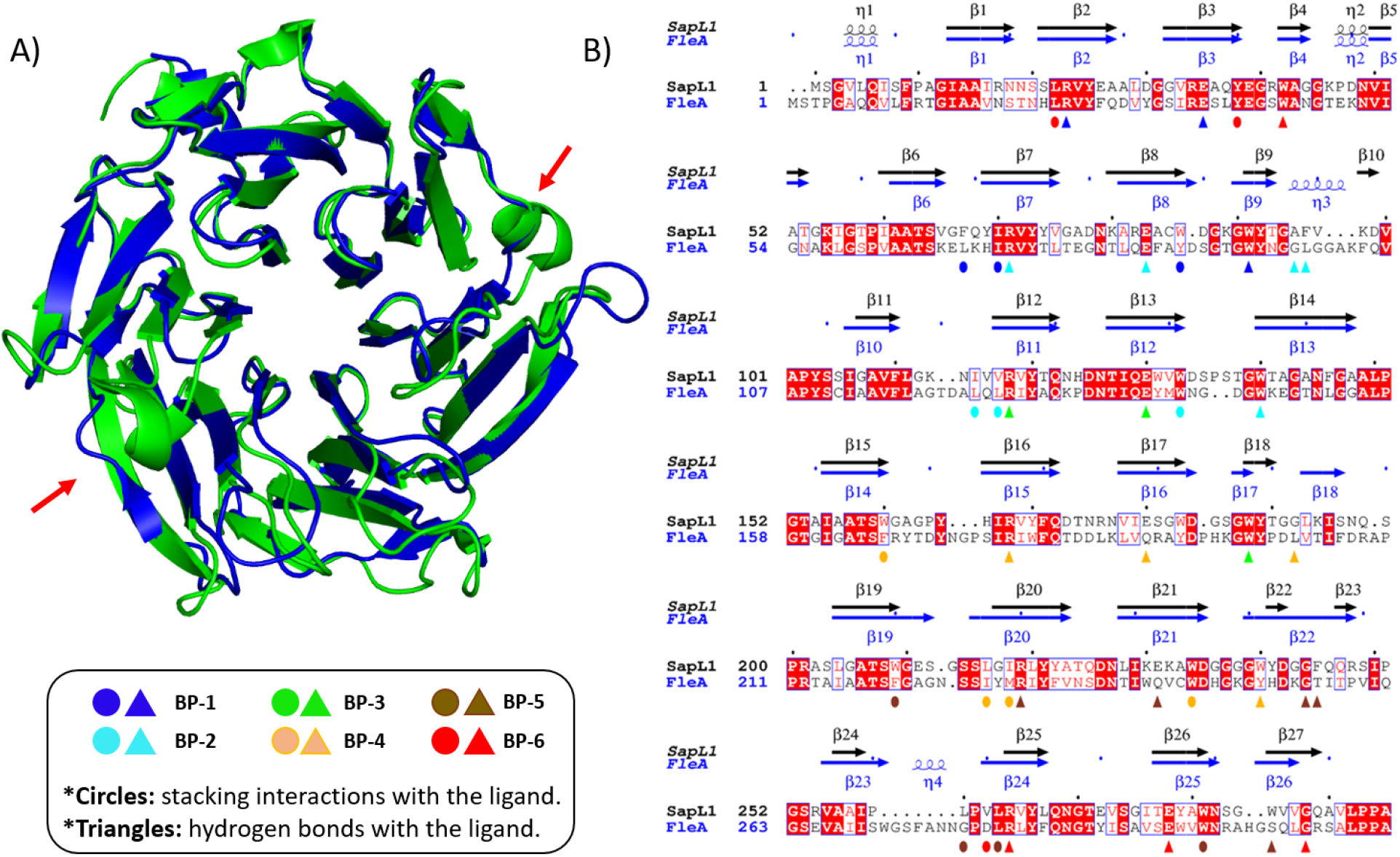
SapL1/FleA comparison. A) Overlay of SapL1 (blue, PDB: 6TRV) and FleA (green, PDB: 4D4U) structures. Red arrows indicate the main differences between structures. B) Sequence alignment of SapL1 and FleA with display of their secondary structure elements. The residues involved in ligand interactions of each pocket are indicated by circles for hydrophobic and stacking interactions and by triangles for hydrogen bonds colored according to the binding pocket: BP-1, blue; BP-2 cyan; BP-3 green; BP-4, orange, BP-5, brown; BP-6, red.

### Protein-ligand interactions

Due to divergence in the tandem repeat sequence forming each blade of the propeller, the six binding sites of SapL1 monomer are not equivalent, but they share important conserved features. Hydrophobic interactions are observed between the C6 of the fucose and at least three residues of the protein (mainly isoleucine, tryptophan/tyrosine and leucine). The O2 and O3 hydroxyls make strong hydrogen bonds with a conserved triad of amino acids consisting of an arginine, a glutamic acid and a tryptophan. In the cases where glycerol was found in the binding pocket, its own hydroxyls mimic the interactions of fucose with these same residues (Fig. 6). It is to be noted that SapL1 binding sites are more conserved than FleA binding sites where a glutamine can replace the glutamic acid and a tyrosine the tryptophan in the triad (Fig. S5).

**Figure 6:**
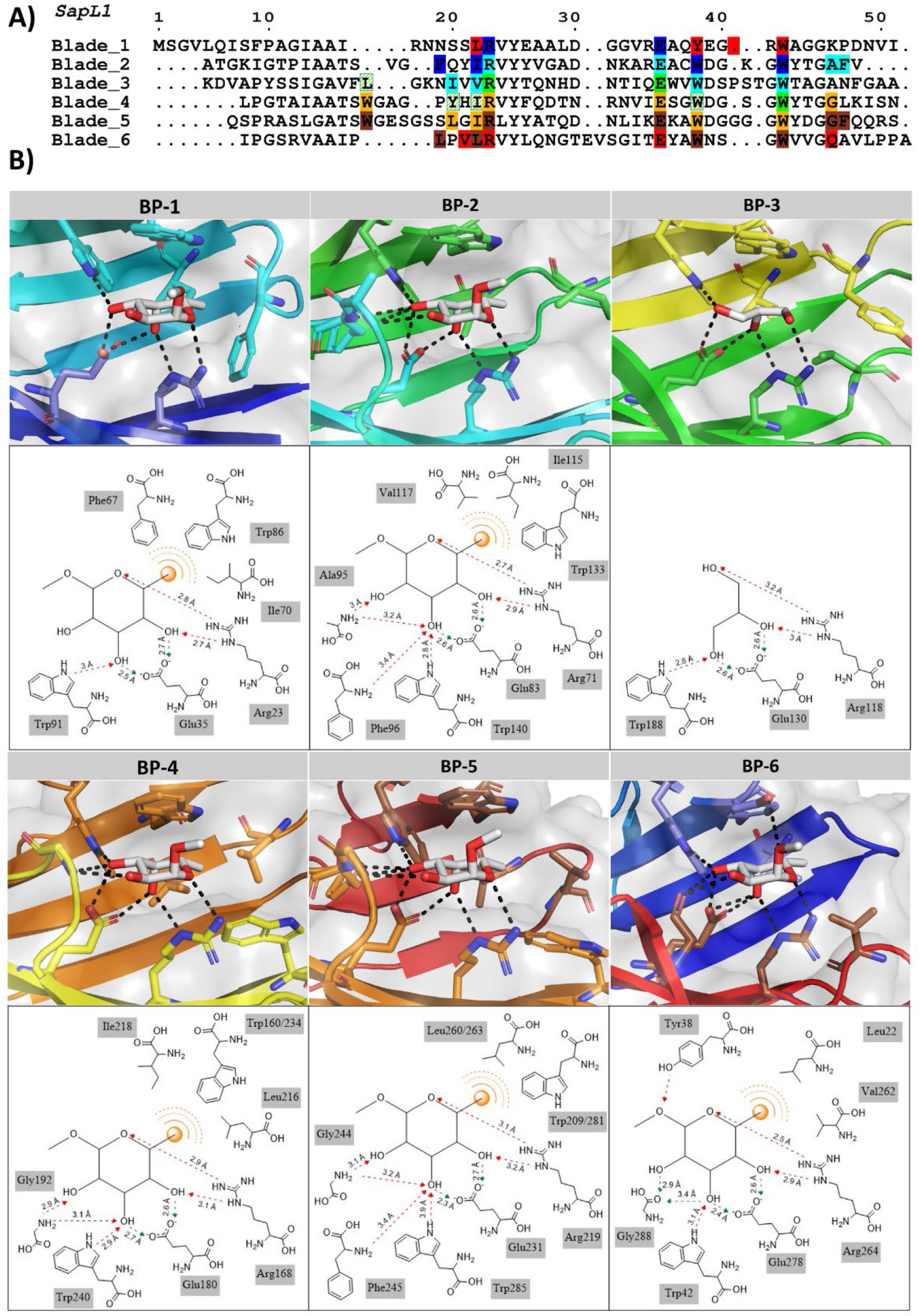
SapL1-ligand interactions. A) The sequence alignment of the 6 blades of SapL1. The residues involved in ligand binding interactions are indicated in solid boxes: BP-1, blue; BP-2 cyan; BP-3 green; BP-4, orange, BP-5, brown; BP-6, red. Striped green boxes indicate the four additional residues expected to be involved in fucose binding within BP-3 that were not visible in the structure since only glycerol was attached to this pocket. B) Zoom on the interactions of ligand with each binding site of SapL1 and their schematic representation.

The O2 hydroxyl seemed to be the most versatile position, since it established interactions with a loop adjacent in four of the six binding sites (BPs 2, 4, 5 and 6). This is particularly interesting since the binding pockets that do not contain this loop (BPs 1 and 3) were mostly occupied by glycerol instead of fucose, indicating that those interactions could be responsible for enhancing the affinity and should be explored for development of inhibitors.

### Binding to epithelial cells

In order to investigate the role of SapL1 in host-pathogen interactions and especially in adhesion, fluorescence microscopy was used. FITC-labelled SapL1 was incubated at two different concentrations with BEAS-2B human bronchial epithelial cells. The microscopy images, obtained for two separate experiments, clearly show binding of the lectin to those cells (Fig. 7). Fluorescence was observed all around the cell surface but also concentrated in some part of the nucleus. A strong fluorescence signal was already observed at the lowest concentration used (5 μg mL^−1^). The binding of SapL1 was inhibited in the presence of its cognate ligand: α-methyl-fucoside (Fig. 7E-G). Some background signal was still observed in the first experiment (Fig. 7E and 7G) whilst it was totally abolished in the duplicate (Fig. 7F and 7H). These data show that SapL1 binding to the cells is dependent on the recognition of fucosylated carbohydrates.

**Figure 7:**
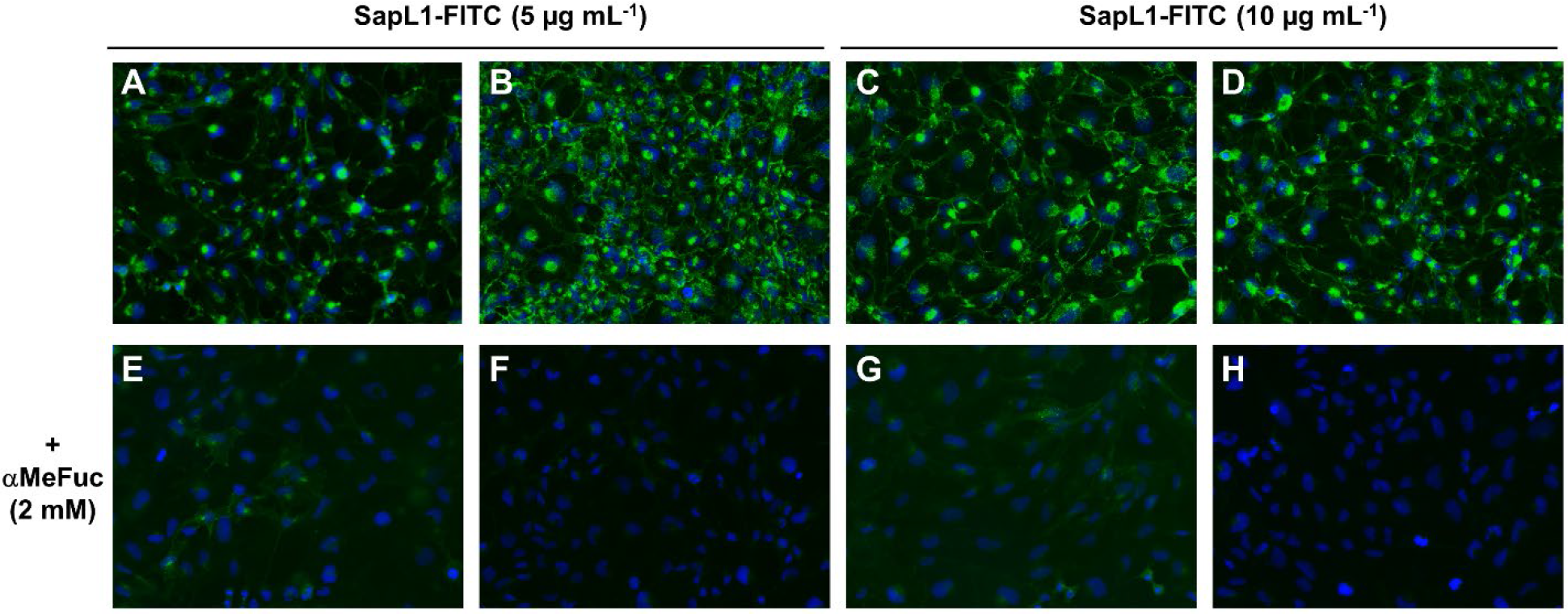
Interaction of *Scedosporium apiospermum* lectin SapL1 with bronchial epithelial cells. A, B, C, D) Representative microscopic images of bronchial epithelial cells (BEAS-2B cell line) treated with SapL1-FITC (green) (5 (A,B) or 10 (C,D) μg mL^−1^) for 1 hour. E, F, G, H) Representative microscopic images of bronchial epithelial treated for 1 hour with SapL1-FITC (green) (5 (E,F) or 10 (G,H) μg mL^−1^) previously preincubated 30 min with 2 mM methyl-α-L-fucopyranoside (αMeFuc). Nuclei were labelled with 4’-6-diamidino-2-phenylindole dihydrochloride (DAPI, blue). Cells were visualized with an Olympus BX43 microscope (magnification ×20).

## Discussion

In comparison with pathogenic bacteria, which have received particular attention for several years, there is a lack of information about the virulence factors and host recognition systems of pathogenic fungi^38^. This hinders the development of new drugs against these microorganisms, while their resistance to current antifungals is progressing quickly^9,21,39^. Therefore, there is a rush in obtaining information that may contribute to the development of new antifungal agents.

In this study, we present the identification, production and characterization of a new carbohydrate-binding protein from the emerging microfungus *S. apiospermum*, whose host-anchoring mechanisms are completely unknown. To the best of our knowledge, this is the first studied lectin for this opportunistic pathogen. SapL1 is homologous to the conidial surface lectin FleA from *A. fumigatus* known to be involved in adhesion to the host glycoconjugates present in bronchial mucus and human lung epithelium^28^. When we analyzed the amino acid sequence deduced from SAPIO_CDS9261, we found at its N-terminus, a peptide sequence of 74 amino acids that was lacking in any other related protein. We believed in a possible error in the annotation of the open reading frame (ORF) of SapL1 since this peptide, predicted to be disordered, cannot be found in other protein and was not found in any other lectin from this family. We failed, however, to amplify the gene from total mRNAs from mycelium but recent transcriptomics data revealed that sapL1 is not usually expressed in mycelium apart in some condition at low pH and in hypercapnia explaining our negative results^40^. By analogy to FleA, it is probably expressed in conidia and new amplification trials should be done on total mRNAs from conidia, but we did not have access to some at that time and there are no transcriptomic data available for this morphological stage to date. When looking at the transcript KEZ40204 for SAPIO_CDS9261, we noticed that the first exon containing the actual initial methionine is only 8 bases long followed by a very long intron of 686 bases which is quite unusual in fungi. After alignment of this transcript sequence from all genome sequences available for *Scedosporium* species, we saw that the proposed initial methionine is not conserved contrary to the one for Met75 (Fig. S6). Whilst waiting for experimental data, this strengthens our hypothesis for a gene misannotation and provides additional information that supports our decision to study the construct of SapL1 comprising only the predicted CRD.

In our initial attempts to produce recombinant SapL1, we observed that the conventional parameters of expression in *E. coli*, resulted in the production of insoluble protein that was mainly found in inclusion bodies. Later, experimental data indicated that the presence of rhamnose or glycerol was required for SapL1 solubilization during bacterial expression and that the production yield was directly proportional to their concentration in the culture medium. We initially considered that the higher production rates of SapL1 could be attributed to the dissociation of the *LacI*-DNA complex caused by an increase in cytosolic potassium concentration. The accumulation of this ion is one of the main adaptive responses of *E. coli* to hypertonic shocks and its concentration has been correlated to the loss of interactions between DNA and proteins in *in vitro* assays^41–43^. However, previous reports have demonstrated that, *in vivo*, the effects of macromolecular crowding caused by plasmolysis (the reduction of intracellular water content as an adaptive response to osmotic shock) on the activity of cytoplasmic proteins is sufficient to buffer the kinetics of association of DNA-protein against changes in [K^+^]^43,44^. Thus, it seems that the higher recovery rates of SapL1 actually were caused by a combination of two main parameters: 1) the leak of genetic repression by dissociation of the *LacI-DNA* complex, that is characteristic of *lac* promoters and 2) a positive effect on the solubility and folding of the protein due to the increased concentration of organic osmoregulators, whose synthesis is induced in *E. coli* as an adaptive response to the external hypertonicity caused here by the addition of rhamnose to the culture medium^41–49^. This hypothesis was later supported by the finding that the addition of glycerol, instead of rhamnose, displays the same effect on production and solubility of SapL1 (data not shown). Plasmolysis may also play an essential role in SapL1 stabilization, since it has been shown that macromolecular crowding has positive effects on protein folding and in some cases it can drive self-association of improper folded proteins into functional oligomers^43^. These findings, and the extensive research carried out to produce SapL1 in its soluble form, provide important insights for the heterologous expression of eukaryotic lectins tending to be produced as insoluble proteins in *E. coli.* Besides, our findings highlight the useful role of organic osmoregulators during heterologous expression of proteins to favour their proper folding and to avoid the formation of inclusion bodies.

We demonstrated that SapL1 is strictly specific for fucosylated carbohydrates and recognized all blood group types present in the glycan array screening. These results are particularly interesting since it has been shown that there is six times more fucose α1,3/4 linked in glycoproteins in the CF airways^31^. This phenomenon is mainly due to an increased expression of the α1,3-fucosyltransferase, which is involved in the synthesis of sialyl-Lewis X and Lewis X determinants attached to bronchial mucins^32^. Therefore, the fact that SapL1 recognizes equally the α1,2 and α1,3/4-linked fucosides may explain the high incidence of scedosporiosis in CF patients. Besides, it correlates with the presence of *SapL1* gene in all the pathogenic strains of *Scedosporium* whose genome has been sequenced to date^33,50,51^. However, there is no evidence suggesting that blood group phenotypes may have an influence in this recognition conversely to previous reports on other pathogens like *Pseudomonas aeruginosa* and *Haemophilus influenzae*^52–54^.

SapL1 adopts a the 6-bladed β-propeller fold and forms dimer leading to two fucose binding surfaces. It belongs to the ProLec6A lectin family whose biological role is still unknown, in particular for the mushroom members^35^. The first member structurally characterized in this lectin family was AAL from the orange peel mushroom *Aleuria aurantia*^55^. It presents differences on both sides of the β-propeller with the microfungal members of this family: SapL1, AOL and FleA mainly at the level of the surface loops. On one side, this leads to a different dimerization interface and the dimer cannot overlay. On the other side, where sugar binding occurs, the differences in size and sequence of the surface loops lead to change in the architecture of the related binding sites and affect fucose binding since AAL has only five and not six functional binding sites^27,37,55^. The specificity and affinity of each one of the six binding pockets of SapL1 have been deeply analyzed to highlight the features that must be explored for the design of efficient inhibitors. Within our analysis, we found that the binding pockets are non-equivalent but they all share the features necessary for fucose recognition. This example of divergence can also be found in the other lectins of this family with FleA exhibiting the greatest differences between pockets known to date^34^. It is remarkable that although SapL1 has a relatively low sequence identity with other members of this family, its specificity and affinity are very similar to those reported for FleA from *A. fumigatus*^36^, AAL from *Aleuria aurantia*^55^and the bacterial lectins BambL and RSL from *Burkholderia ambifaria* and *Ralstonia solanacearum,* respectively^56,57^.

Finally, fluorescence microscopy experiments allowed us to get preliminary results on SapL1 function. The lectin is able to recognize and to bind to bronchial epithelial cells, and we demonstrated that this binding is dependent on the recognition of fucosylated glycoconjugates at the surface of the target cells as it is abolished in the presence of fucose. These results corroborate a role of SapL1 in mediating the recognition and adhesion of *S. apiospermum* to host cells as found for other lectins from pathogenic microorganisms and in particular for its homolog FleA^22,27^. Localisation of the lectin in the microfungus would also help to better understand its role but no SapL1 antibodies are yet available. Together with the crystallographic data, our study shows that the structure and function of lectins belonging to the 6-bladed β-propeller fucose specific family recently named PropLec6A are highly conserved, settling the possibility for development of a broad-spectrum therapy^35^.

Overall, our research has revealed the first insights about the recognition of fucosylated human glycoconjugates by *S. apiospermum* lectin SapL1 and contributes to the general understanding of the host-binding process during the early stages of infection. The lectin is found in all *Scedosporium* strains sequenced. The detailed information exposed here places SapL1 as a promising target for treatment of *Scedosporium* infections although it is clear that more functional data are still required. It will be of great value to guide the development of antiadhesive glycodrugs against this pathogen.

## Methods

### Production

The coding sequence for SapL1 (75-369) was optimized for expression in *Escherichia coli* and ordered at Eurofins Genomics (Ebersberg, Germany) before cloning into the expression vectors pET-TEV^58^, pET32-TEV^36^ and pProNde. pProNde is a homemade vector where the NcoI restriction site of pProEx HTb (EMBL, Heidelberg) has been replaced by a NdeI restriction site by PCR. Then, plasmids were introduced into *E. coli* strains by thermal shock at 42°C and different expression parameters were assayed (Table S1) until soluble expression was achieved. Finally, SapL1 was expressed using pProNde vector in *E. coli* TRX(DE3) strain with the cells grown at 37°C and 160 rpm until an OD_600_ of 0.4. The temperature was then lowered to 16°C and when OD_600nm_ reached 0.8, the induction was carried out overnight by the addition 0.05 mM of IPTG and 1% L-rhamnose.

### Purification

Cells were subsequently harvested by centrifugation at 5000 g for 10 min and resuspended in buffer A (50 mM Tris-HCl, 500 mM NaCl, pH 8.5). After addition of 1 μl of Denarase (C-LEcta GmbH, Leipzig, Germany) and moderate agitation during 30 minutes at room temperature, cells were disrupted at 1.9 kbar using a cell disruptor (Constant Ltd Systems, UK). Cell debris were removed by centrifugation at 22000 g for 30 minutes. The supernatant was filtered using 0.45 micrometer membranes (PES, ClearLine) and the protein purification was carried out by IMAC using 1 mL His-Trap FF columns (GE Healthcare Life Sciences) and a NGC chromatography system (Bio-Rad). After loading the supernatant, the column was rinsed thoroughly with buffer A until stabilization of the baseline. Bound proteins were eluted through the addition of buffer B (50 mM Tris-HCl, 500 mM NaCl, 500 mM imidazole, pH 8.5) in an 0-500 mM imidazole gradient over 20 mL. Fractions containing SapL1 were pooled and concentrated by ultrafiltration (Pall, 10kDa cut-off) prior buffer exchange to 50 mM Tris-HCl, 100 mM NaCl, pH 8.5 using PD10 desalting columns (GE Healthcare Life Sciences). Then, the fusion was cleaved off overnight at 19°C using the TEV protease produced in the lab in 1:50 ratio and addition of 0.5 mM EDTA and 0.25 mM TCEP. The sample was loaded on the His-trap column to separate the cleaved protein collected in the flow-through from the TEV protease and potential uncleaved sample retained and eluted with imidazole. Sapl1 containing fractions were concentrated by centrifugation to the desired concentration.

### Size exclusion chromatography

Size exclusion chromatography (SEC) was performed using a High-Resolution ENrich SEC 70 column (Bio-Rad) on NGC™ chromatography system (Bio-Rad). Column was equilibrated with 50 mL of buffer D (20 mM MES, 100 mM NaCl, pH 6.5) and 200 μl of sample at 10 mg mL^−1^ were injected into the system followed by 40 mL isocratic elution on buffer D supplemented with 20 mM α-methyl fucoside, 0.1% L-rhamnose, 5 M NaCl, 2 M NaCl or 500 mM NaCl, according to the experiment. Fractions were monitored by the absorbance at 280 nm and 0.5 mL fractions were collected in the resolving region of the column.

### Dynamic light scattering (DLS)

DLS analyses were performed using a Zetasizer Nano ZS (Malvern Panalytical) with a 40 μl quartz cuvette. Measurements were performed in triplicate on protein sample at 1 mg·mL^−1^ in buffer C (50 mM Tris-HCl, 100 mM NaCl, pH 8.5) after centrifugation.

### Thermal Shift Assay (TSA)

The thermal stability of SapL1 was analyzed by TSA with the MiniOpticon real-time PCR system (Bio-Rad). Prior assay, buffer stocks at 100 mM and a mixture containing 70 μl of SapL1 at 1 mg mL^−1^, 7 μl of 500x Sypro Orange (Merk Sigma-Aldrich,) and 63 μl of ultrapure H_2_O were prepared. Then, 7.5 μl of H_2_O, 12.5 μl of the corresponding buffer and 5 μl of the protein/Sypro mixture were mixed in 96-well PCR microplates. The heat exchange test was then carried out from 20°C to 100°C with a heating rate of 1°C·min^−1^. Fluorescence intensity was measured with Ex/Em: 490/530 nm and the data processing was performed with the CFX Manager software.

### Isothermal Titration Calorimetry (ITC)

Experiments were performed using a Microcal ITC200 calorimeter (Malvern Panalytical) with 40 μl of L-fucose 5 mM in the syringe and 200 μl of protein 0.05 mM in the sample cell. Both, the protein and sugars were dissolved in a buffer composed of 20 mM Tris HCl pH 8.0 and 100 mM NaCl. A total of 26 injections of 1.5 μL of ligand were added to the sample cell at intervals of 180 s while stirring at 850 rpm. Experimental data were adjusted to a theoretical titration curve by the Origin ITC Analysis software. All experiments were performed at least by duplicated and the stoichiometry was fixed to 1.

### Hemagglutination assay

Agglutination test was performed with fresh rabbit erythrocytes (bioMérieux, Lyon) in U 96-well plates (Nalgene). For the test, 150 mM NaCl was used as a negative control and 1 mg mL^−1^ LecB of *P. aeruginosa* as a positive control. 50 μl of sample was prepared at 0.1 mg·mL^−1^ and submitted to serial double dilutions. 50 μl of rabbit erythrocytes 3% were added to each well prior incubation of the plate at room temperature. After 2 hours, the result of the experiment was evaluated and agglutination activity was calculated according to the dilution of the protein.

### Glycan arrays

Protein was labeled with Fluorescein Isothiocyanate (FITC, Merk, Sigma-Aldrich) according to the supplier’s instructions with slight modifications. Briefly two milligrams of protein were dissolved in 1 mL of buffer E (100 mM Na_2_CO_3_, 100 mM NaCl, pH 9); then, 40 μl of FITC at 1 mg·mL^−1^, previously dissolved in DMSO, were gradually added to the protein solution and the mixture was gently stirred at room temperature overnight. Next day, the solution was supplemented with NH_4_Cl to a final concentration of 50 mM and free FITC was removed using PD10 column with PBS as mobile phase. Protein concentration was determined at *A*_280_ and FITC at *A*_490_ using a NanoDrop 200 (Thermo Scientific) and Fluorescein/Protein molar ratio (F/P) was estimated by the following formula:

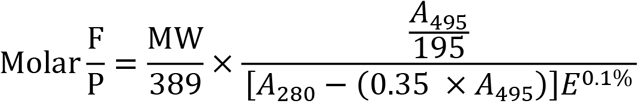

Where MW is the molecular weight of the protein, 389 is the molecular weight of FITC, 195 is the absorption E 0.1% of bound FITC at 490 nm at pH 13.0, (0.35 X *A*495) is the correction factor due to the absorbance of FITC at 280 nm, and E 0.1% is the absorption at 280 nm of a protein at 1.0 mg·mL^−1^, Being an ideal F/P should be 0.3 > 1.

Labeled lectin was sent to the Consortium for Functional Glycomics (CFG; Boston, MA, USA) and its binding properties were assayed at 5 and 50 μg·mL^−1^ on a “Mammalian Glycan Array version 5.4” which contain 585 glycans in replicates of 6. The highest and lowest signal of each set of replicates were eliminated and the average of the remaining data was normalized to the percentages of the highest RFU value for each analysis. Finally, the percentages for each glycan were averaged at different lectin concentrations.

### Crystallization and data collection

Crystal screening was performed using the hanging-drop vapour diffusion technique by mixing equal volumes of pure protein at 5 mg·mL^−1^ and precipitant solutions from commercial screenings of Molecular Dimensions (Newmarket, UK). 2 μl drops were incubated at 19°C until crystals appeared. A subsequent optimization of positive conditions for SapL1 crystallization was carried out and crystals suitable for X-ray diffraction analysis were obtained under solution containing 100 mM Bicine pH 8.5, 1.5 M ammonium sulfate (NH_4_)_2_SO_4_ and 12% v/v glycerol. Crystals were soaked in mother liquor supplemented with 10% (v/v) glycerol, prior to flash cooling in liquid nitrogen. Data collection was performed on PX1 beamline at SOLEIL Synchrotron (Saint Aubin, France) using an Eiger2 X 9M pixel detector (Dectris Ltd, Switzerland).

### Structure determination

Data were processed using XDS^59^ software and were converted to structure factors using the CCP4 program package v.6.1^60^, with 5% of the data reserved for Rfree calculation. The structure was determined using the molecular-replacement method with Phaser v.2.5^61^, using the structure of FleA dimer (PDB entry 4D4U^34^) as starting model. Model refinement was performed using REFMAC 5.8^62^ alternated with manual model building in Coot v.0.7^63^. Sugar residues and other compounds that were present were placed manually using Coot and validated using Privateer^64^. The final model has been validated and deposited in the PDB Database with accession number 6TRV.

### Cell culture and fluorescence microscopy

Human bronchial epithelial cells (BEAS-2B cell line) were maintained and serially passaged in F-12 culture medium supplemented with 10% fetal calf serum (FCS), 1% penicillin and streptomycin, and 10 mM HEPES in 75-cm^2^ culture flasks. For microscopy experiments, BEAS-2B cells were grown to confluency on coverslips (precision cover glasses thickness No. 1.5H; Marienfeld, Lauda-Königshofen, Germany). To test interaction of SapL1 lectin with epithelial cells, BEAS-2B cells were incubated with different concentrations of FITC-lectin (5 or 10 μg mL^−1^ in F-12) for 1 hour at 37°C. To validate the specificity of this interaction, SapL1-FITC was co-incubated in presence of 2 mM methyl-α-L-fucopyranoside for 30 min at 37°C and then added to the cells for 1 hour. Supernatant was then removed, cells washed 3 times with F-12, once with PBS and then fixed with 4% paraformaldehyde for 15 min. After 3 washes with PBS, nuclei were stained with 4′-6-diamidino-2-phenylindole dihydrochloride (DAPI, 1:1000 in PBS) for 5 min. Coverslip were mounted with Prolong Glass Antifade Mountant (Invitrogen) on Superfrost glass slides (Thermo Fisher Scientific). Images were acquired with an upright Olympus BX43 microscope.

### Figures

Figures were created using PyMOL Version 1.8.4 (Schrödinger), ChemDraw Version 15, ESPript Version 3.0^65^ and PowerPoint 16.

## Supporting information

Supplemental images and table

## Acknowledgments

We would like to thank to Valérie Chazalet and Emilie Gillon for their technical assistance. We are grateful to the Consortium for Functional Glycomics for providing the glycan array resource. We also would like to thank for access to the beamlines BM30A-FIP at the European Synchrotron Radiation Facility (ESRF), Grenoble, France where initial tests were performed and Proxima 1 at SOLEIL Synchrotron, Saint Aubin, France (proposal number 20170827), where final data were collected. Thanks to our local contacts Jean-Luc Ferrer, Serena Sirigu, and Pierre Legrand for their assistance and technical support.

## Author Contributions

D.M.A. performed cloning, protein expression, protein purification, protein labelling, hemaglutination, isothermal microcalorimetry measurements. V.B. performed the fluorescence microscopy experiments. D.M.A. participated in all steps of protein crystallography and data analysis, wrote the original draft of the manuscript and prepared all figures. J-P.B. provided the genome data for the bioinformatics studies and contributed to revise the manuscript. R.J.P. participated in the supervision of D.M.A, funding acquisition and revision of the manuscript. A.V. administered the project, conceived the design of the study, obtained funding, evaluated the results, contributed to data analysis, supervised D.M.A., helps in the preparation of the manuscript and contributed to revise the manuscript and the figures. All authors have read and agreed to the published version of the manuscript.

## Accession numbers

The structure of SapL1 was deposited as PDB ID: 6TRV.

