## Supplemental images and table for "Biochemical and structural studies of target lectin SapL1 from the emerging opportunistic microfungus *Scedosporium apiospermum*"

* Corresponding author

**S1 Table.** SapL1 expression conditions assayed.

<table>
<tr><th>Strain</th><th>vector</th><th>Media</th><th>Inductor</th><th>Inductor concentration</th><th>$A_{600}$ at induction</th><th>Induction temperature</th><th>Expression length</th></tr>
<tr><td rowspan="20">BL21(DE3)</td><td>pET-TEV</td><td rowspan="63">LB</td><td rowspan="39">IPTG</td><td rowspan="3">100 µM</td><td rowspan="18">0.8</td><td rowspan="2">16°C</td><td rowspan="4">Overnight</td></tr>
<tr><td rowspan="6">pProNde</td></tr>
<tr><td rowspan="5">25°C</td></tr>
<tr><td>250 µM</td></tr>
<tr><td>100 µM</td><td rowspan="3">36 h</td></tr>
<tr><td>250 µM</td></tr>
<tr><td>150 µM</td></tr>
<tr><td rowspan="13">pET32-TEV</td><td rowspan="7">100 µM</td><td rowspan="38">16°C</td><td>1 h</td></tr>
<tr><td>2 h</td></tr>
<tr><td>3 h</td></tr>
<tr><td>4 h</td></tr>
<tr><td>5 h</td></tr>
<tr><td>6 h</td></tr>
<tr><td rowspan="16">Overnight</td></tr>
<tr><td>50 µM</td></tr>
<tr><td>25 µM</td></tr>
<tr><td>10 µM</td></tr>
<tr><td>5 µM</td></tr>
<tr><td rowspan="3">100 µM</td><td>2</td></tr>
<tr><td>1.5</td></tr>
<tr><td rowspan="11">Tuner™(DE3)</td><td rowspan="6">pProNde</td><td rowspan="2">2</td></tr>
<tr><td rowspan="2">250 µM</td></tr>
<tr><td>2.5</td></tr>
<tr><td>100 µM</td><td>0.8</td></tr>
<tr><td>250 µM</td><td rowspan="4">0.8</td></tr>
<tr><td>10 µM</td></tr>
<tr><td rowspan="5">pET32-TEV</td><td>100 µM</td></tr>
<tr><td>250 µM</td></tr>
<tr><td rowspan="3">100 µM</td><td>1</td></tr>
<tr><td rowspan="2">0.8</td><td>36 h</td></tr>
<tr><td rowspan="37">Overnight</td></tr>
<tr><td rowspan="4">BL21Star(DE3) pLysS</td><td rowspan="4">pProNde</td><td>25 µM</td><td>2</td></tr>
<tr><td rowspan="4">100 µM</td><td>2</td></tr>
<tr><td>1.7</td></tr>
<tr><td rowspan="15">0.8</td></tr>
<tr><td rowspan="2">Rosetta™(DE3) pLysS</td><td rowspan="4">pET32-TEV</td></tr>
<tr><td>25 µM</td></tr>
<tr><td rowspan="2">Rosetta-gami 2 (DE3)</td><td>100 µM</td></tr>
<tr><td>10 µM</td></tr>
<tr><td rowspan="4">KRX (DE3)</td><td rowspan="4">pProNde</td><td rowspan="19">Rh</td><td>0.1%</td></tr>
<tr><td>0.5%</td></tr>
<tr><td>1%</td></tr>
<tr><td>2.5%</td></tr>
<tr><td rowspan="17">BL21trxB (DE3)</td><td rowspan="2">pET32-TEV</td><td>0.1%</td></tr>
<tr><td>0.5%</td></tr>
<tr><td rowspan="16">pProNde</td><td>0.1%</td><td rowspan="4">20°C</td></tr>
<tr><td>0.15%</td></tr>
<tr><td>0.2%</td></tr>
<tr><td>0.25%</td></tr>
<tr><td rowspan="2">0.1%</td><td>2</td><td rowspan="18">16°C</td></tr>
<tr><td rowspan="17">0.8</td></tr>
<tr><td>0.15%</td></tr>
<tr><td>0.2%</td></tr>
<tr><td>0.25%</td></tr>
<tr><td>0.5%</td></tr>
<tr><td>0.75%</td></tr>
<tr><td>1%</td></tr>
<tr><td>2%</td></tr>
<tr><td rowspan="2">IPTG/Rh</td><td>50 µM/ 1%</td></tr>
<tr><td>100 µM/ 1%</td></tr>
<tr><td>BL21(DE3)</td><td>Rh</td><td>1%</td></tr>
<tr><td rowspan="4">BL21trxB (DE3)</td><td rowspan="2">pET32-TEV</td><td rowspan="2">IPTG</td><td>100 µM</td></tr>
<tr><td>50 µM</td></tr>
<tr><td rowspan="4">pProNde</td><td>SuperiorLB</td><td>IPTG/Rh</td><td rowspan="4">50 µM/ 1%</td></tr>
<tr><td rowspan="3">LB</td><td>IPTG/Gly</td></tr>
<tr><td rowspan="2">BL21(DE3)</td><td>IPTG/Rh</td></tr>
<tr><td>IPTG/Gly</td></tr>
</table>

```
SAPIO_CDS9261 MVDLGSMTEATTFLIKKYRMIFAEITKGDRLRGGEKSRRRQLEKYGLISQTQDTSSAKSNYSSESLSPQNQQLAMSGVLQ 80
FleA          ------------------------------------------------------------------------MSTPGAQQ 6
                                                                                      ::  *. *

SAPIO_CDS9261 ISFPAGIAAIRNNSSLRVYEAALDGGVREAQYEGRWAGGKPDNVIATGKIGTPIAATSVGFQYIRVYYVGADNKAREACW 160
FleA          VLFRTGIAAVNSTNHLRVYFQDVYGSIRESLYEGSWANGTEKNVIGNAKLGSPVAATSKELKHIRVYTLTEGNTLQEFAY 88
              : * :****:.... ****   : *.:**: *** **.*. .***...*:*:*:****  :::**** :  .*. :* .:

SAPIO_CDS9261 -DGKGWYTGAFV---KDVAPYSSIGAVFLGK--NIVVRVYTQNHDNTIQEWVWDSPSTGWTAGANFGAALPGTAIAATSW 234
FleA          DSGTGWYNGGLGGAKFQVAPYSCIAAVFLAGTDALQLRIYAQKPDNTIQEYMWNG--DGWKEGTNLGGALPGTGIGATSF 166
               .*.***.*.:     :*****.*.****.    : :*:*:*: ******::*:.   **. *:*:*.*****.*.***:

SAPIO_CDS9261 GAGPY---HIRVYFQDTNRNVIESGWD-GSGWYTGGLKISN-QSPRASLGATSWGESGSSLGIRLYYATQDNLIKEKAWD 309
FleA          RYTDYNGPSIRIWFQTDDLKLVQRAYDPHKGWYPDLVTIFDRAPPRTAIAATSFGAGNSSIYMRIYFVNSDNTIWQVCWD 246
                  *    **::**  : :::: .:*  .*** . :.* :   **:::.***:* ..** ::*:*:...** * : .**

SAPIO_CDS9261 GGGGWYDGGFQQRSIPGSRVAAIP-------LPVLRVYLQNGTEVSGITEYAWNSG--WVVGQAVLPPA   342
FleA          HGKGYHDKGTITPVIQGSEVAIISWGSFANNGPDLRLYFQNGTYISAVSEWVWNRAHGSQLGRSALPPA   286
               * *::* *     * **.** *         * **:*:**** :*.::*:.** .    :*::.****
```

**Figure S1. Identification of SapL1.** Alignment of the SAPIO_CDS9261 and FleA sequences. In red, the first 74 residues that were removed from the sequence of SapL1 synthetic gene. Black arrow indicates the first methionine of the recombinant SapL1

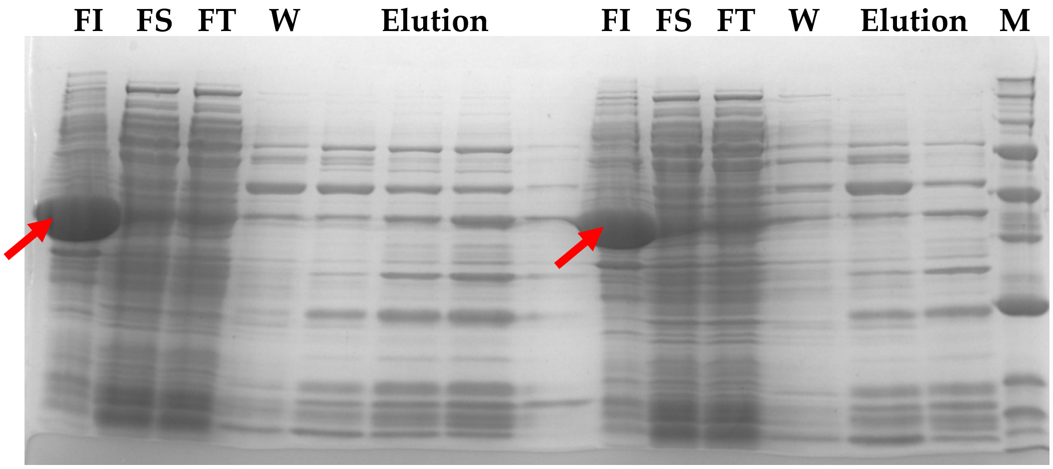


**Figure S2. SapL1 production.** SDS-PAGE of fractions collected from SapL1 purification using pET32-TEV vector. From right to left: insoluble fraction(IF), Soluble fraction (SF), flow-through (FT), wash (W) and elution fractions, respectively ending with the molecular weight marker (M). The left half of the figure shows the purification fractions in an experiment performed with 0.025 mM of IPTG as inductor of SapL1 expression, while the right half of the gel shows the same distribution of samples from an experiment performed with 0.05 mM of IPTG. Molecular weight marker bands: 250, 150,100,**75**, **50**, 37, **25**, 20, 15, 10 kDa from top to bottom. Red arrow indicates inclusion bodies from insoluble fractions.

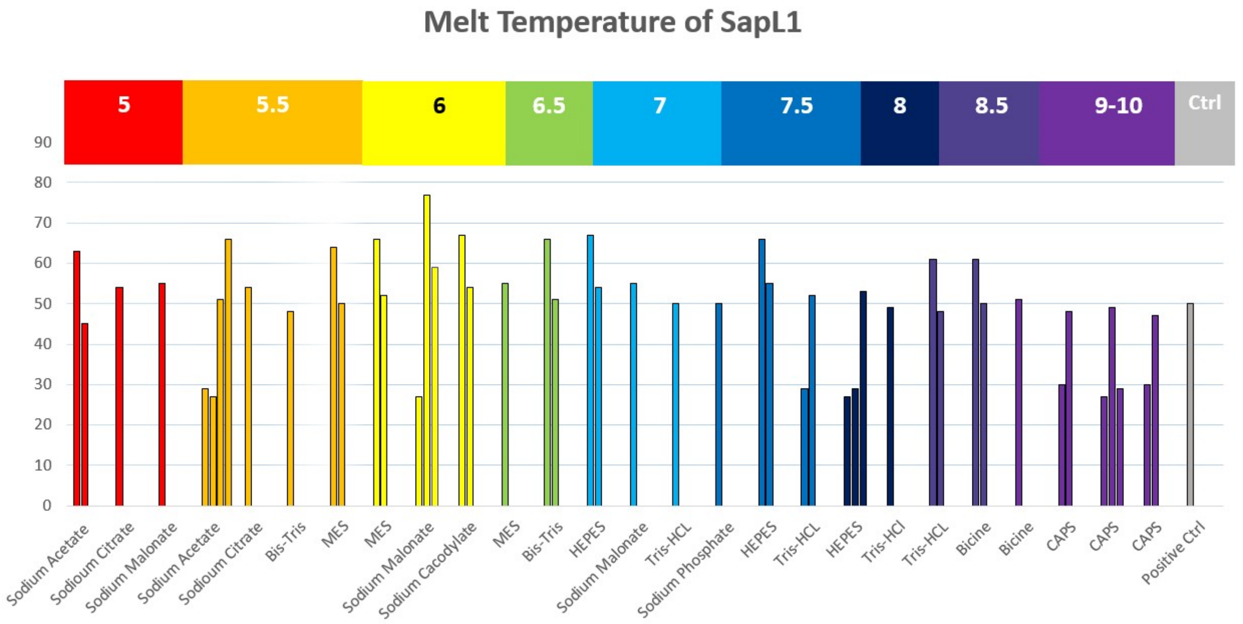


**Figure S3. SapL1 thermal Stability**. Melting temperatures of SapL1 obtained through the Thermal shift assay (TSA). A temperature gradient from 20 to 100°C in was applied under 26 different buffer conditions with a pH range from 5 to 10.

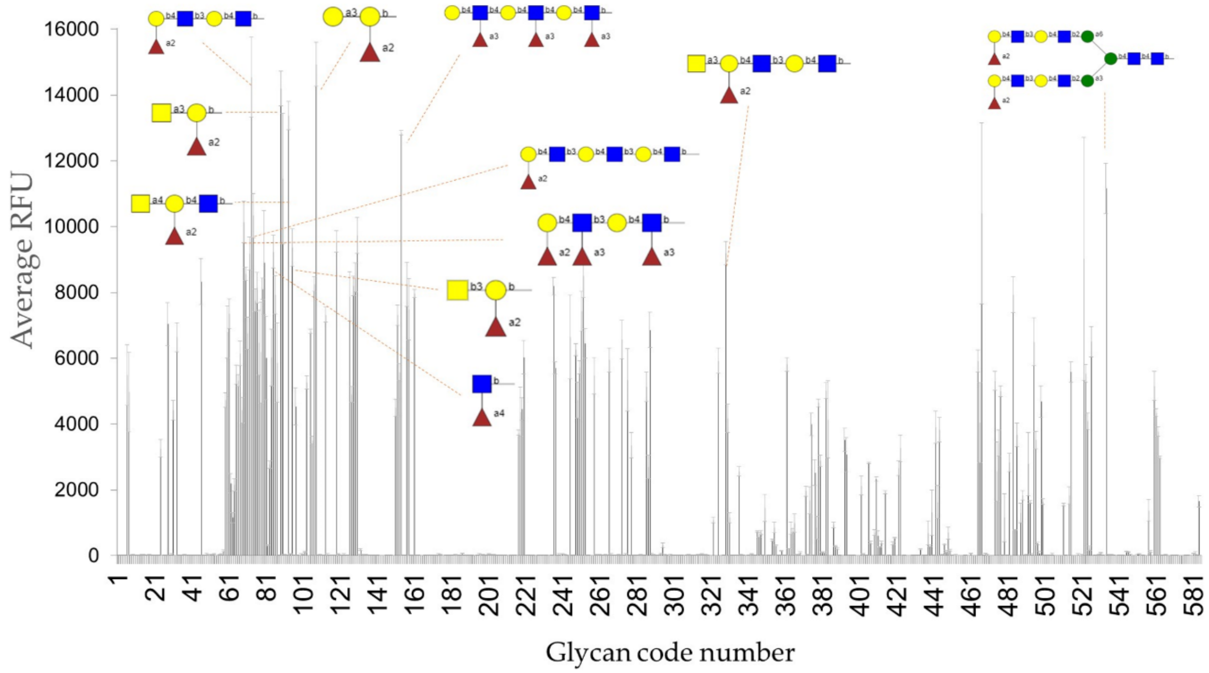


**Figure S4. Glycan array signals.** Relative Fluorescent Units (RFU) plot of the glycan array matrix with SapL1 at 50 μg·ml$^{-1}$. The structures of the ten best binders are represented linked to their respective signals.

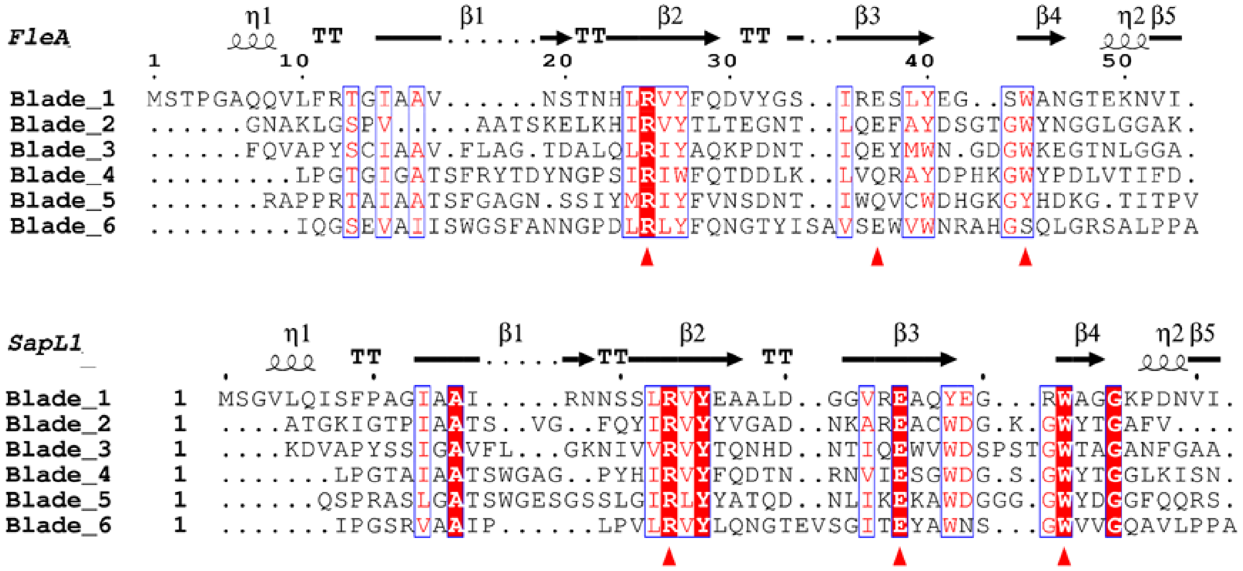


**Figure S5. Conservation of blades in SapL1 and FleA structures**. The conserved (SapL1) and semi conserved (FleA) triads of amino acids involved in ligand binding by hydrogen bonds are indicated with red triangles.

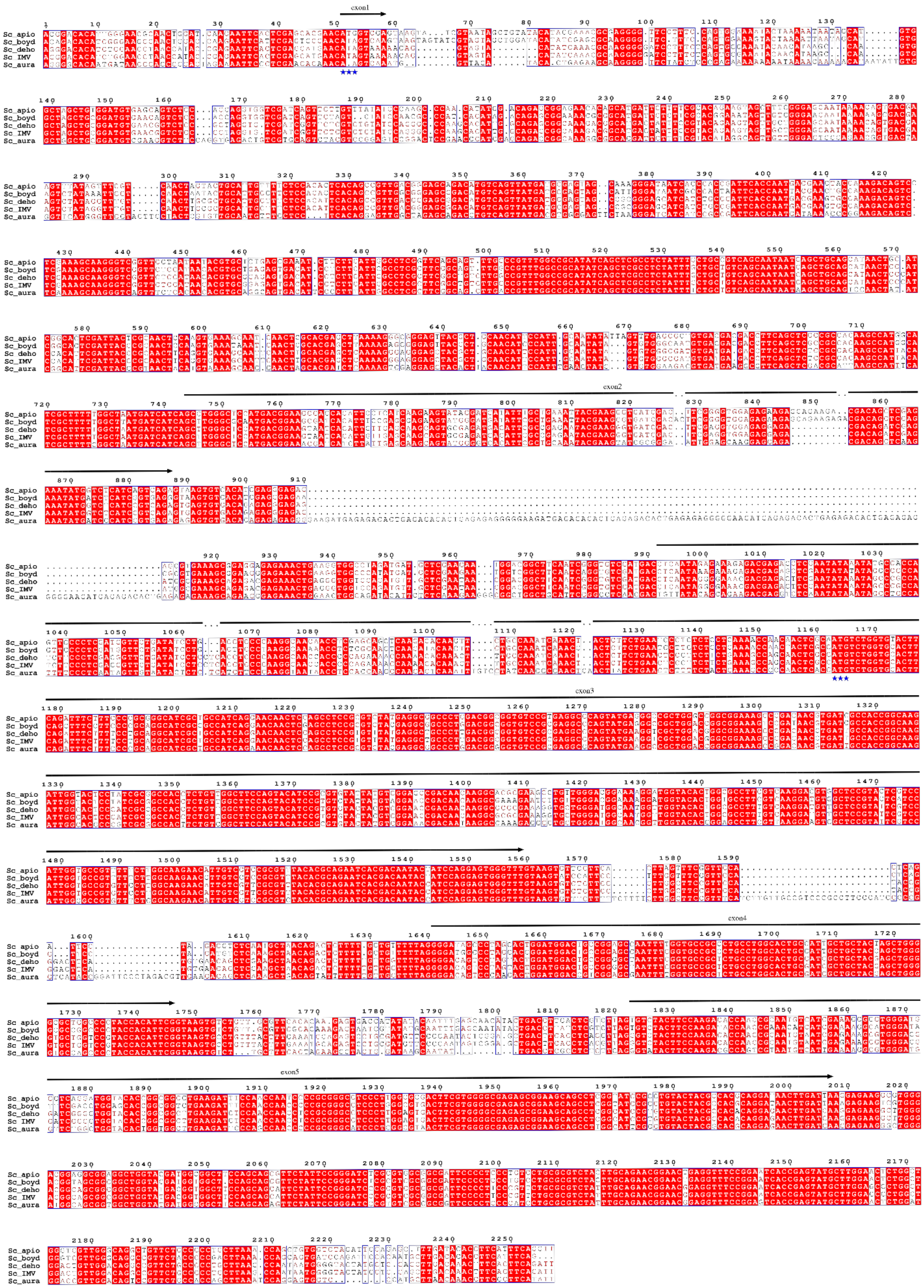
exon1
exon2
exon3
exon4
exon5
Sc_apio
Sc_boyd
Sc_deho
Sc_IMV
Sc_aura

**Figure S6. Alignment of the transcript containing *sapL1* from all Scedosporium species with sequenced genomes**. Sc_apio: *S. apiospermum* SEQ_SAPIO_0132:1935229:1937485[1], Sc_boyd: *S. boydii* strain IHEM 23826 contig 288[2], Sc_deho: *S. dehoogii* strain 120008799-01/4 contig 122, SC_IMV, S. sp. IMV 00882 and Sc_aura: *S. aurantiacum* strain WM 09.24 scaffold-55[3]. Sequence were obtained from NCBI genome page. Prediction exons are depicted by arrows at the top and the two possible initial codons by blue stars at the bottom (positions 51 and 1163). Alignment done using Multalin[4] and figure drawn in ESPript 3.0[5].
